# High methane flux in a tropical peatland post-fire is linked to homogenous selection of diverse methanogenic archaea

**DOI:** 10.1101/2023.04.10.536191

**Authors:** Aditya Bandla, Hasan Akhtar, Massimo Lupascu, Rahayu Sukmaria Sukri, Sanjay Swarup

## Abstract

Tropical peatlands in South-East Asia are some of the most carbon dense ecosystems in the world. Recurrent wildfires in repurposed peatlands release massive amounts of carbon and other greenhouse gases, strongly alter peat geochemistry and physicochemical conditions. However, little is known about the impact of fire on peat microbiome composition, microbial guilds contributing to greenhouse gas emissions, and their predictability based on environmental conditions. Here, we address this gap by studying peat microbiomes from fire-affected and intact areas of a tropical peatland in Brunei using high-throughput sequencing and ecological process modelling at the community and clade levels. We show that fire disrupts depth-stratification of peat microbiomes with the strongest effects observed at 1m below the surface. The enrichment of specific taxa and methanogenic archaea at such depths suggests an adaptation to low-energy conditions post-fire. Finally, fire shifts archaeal community composition and clades containing abundant methanogens in a homogeneous manner that can be predicted from environmental conditions and functional traits. Together, our findings provide a biological basis for earlier work which reported elevated methane flux 2-3 years post-fire and show that such changes follow predictable trajectories with important implications for post-fire microbiome forecasting and ecosystem recovery efforts.

## Introduction

Tropical peat swamp forests store 80-105 Gt carbon (approx. 20% of global peatland carbon) (1, 2) and are highly resistant to wildfires in their natural state (3, 4). Deforestation, drainage-based agriculture coupled with climate extremes have resulted in a significant proportion of peatlands in South-East Asia being subject to frequent and severe disturbances such as wildfires (5). Recurrent wildfires strongly alter peat geochemistry (6, 7), hydrology (8), carbon dynamics (7) and significantly limit forest recovery (9). In contrast to the effects of flaming wildfires on mineral soil geochemistry, which are typically limited to the surface, smouldering peat fires burn deep into the peat and cause wide-ranging changes in peat geochemistry far below the surface (10, 11). Such fire-induced changes in peat geochemistry have in turn been linked to shifts in carbon dynamics which can persist for several years post-fire (7). Although carbon dynamics and forest recovery are closely linked to belowground microbiomes, little is known about the impact of post-fire changes in peat geochemistry and porewater physiochemical conditions on peat microbiomes.

Peat microbiomes regulate carbon processing (12), greenhouse gas emissions (13, 14), nutrient dynamics (15) and plant growth (16, 17). Similar to the effects of wildfires on mineral soil microbiomes (18, 19), a drastic reduction in bacterial abundance at the peat surface (approx. 99%) was reported immediately after fire in a tropical peatland with little to no change observed three years post-fire. In addition to changes at the surface, a two-third decrease in bacterial abundance was observed in sub-surface peat (30-50 cm) highlighting the effects of peat fires deep below the surface (20). Fires in mineral soil ecosystems reduce microbiome diversity (19, 21, 22), strongly shift composition (22, 23, 24), and alter the abundance of key functional groups involved in carbon turnover (22, 25, 26, 27), however, this is yet to be established in fire-affected tropical peatlands.

In addition to effects on microbiome diversity and composition, fires affect the predictability of microbiome compositional shifts by altering the relative balance between deterministic and stochastic assembly processes. A limited number of studies on mineral soil ecosystems have shown that the predictability of microbiome shifts post-fire depends on time since fire (27, 28, 29). An increase in stochastic processes such as dispersal and ecological drift was observed for a brief period (4-16 weeks) post-fire likely due to limited competition and weak environmental cues (28). Following this initial period, deterministic processes such as homogeneous selection have been shown to increase 1-4 years post-fire (30) during which fire-induced changes in soil chemistry and aboveground vegetation act as strong environmental filters on microbiome composition. Severe, recurrent wildfires homogenize environmental conditions and result in convergent or homogeneous selection (22) while those of low severity lead to divergent or heterogeneous selection due to variable effects on the environment (31). Finally, recovery from fire on longer time scales (>25 years) is characterised by an increase in stochastic processes as environmental conditions begin to normalize. Although such studies provide valuable insights into compositional predictability at the community-level, they do not provide information on the predictability of individual clades which may differ in their response to fires. Phylogenetic-bin based null modelling approaches can provide insights into compositional predictability both at the community and clade levels (32). A recent study showed that individual clades whose members occupy similar ecological niches do indeed respond differently to fires and consequently are shaped by different assembly processes (27). Such approaches thus have the potential to provide an understanding of assembly processes acting on key functional groups such as those that turnover carbon.

Here we link altered peat physicochemical conditions and high methane emissions from a fire-affected tropical peatland to shifts in peat microbiome composition, potential function, and compositional predictability using high-throughput 16S rRNA marker gene sequencing and ecological process modelling. Our study builds upon previous work (7) which showed that peat porewater physicochemical conditions, peat geochemistry and carbon dynamics (elevated methane flux) remained strongly altered 2-3 years post-fire in a tropical peatland in Brunei. Thus, we hypothesised that fires will shift microbiomes and important carbon processing guilds such as methanogens in a predictable manner with effects persisting for several years post-fire. We found that fire significantly shifts microbiome composition and increases the abundance of diverse methanogenic archaea. Post-fire physicochemical conditions impose strong, consistent selective pressures on archaeal communities and methanogens resulting in highly homogenised communities. Overall, this study furthers our understanding of the linkages between shifts in tropical peat microbiomes and altered carbon dynamics post-fire.

## Methods

### Sample collection and measurement of physicochemical conditions

Peat samples were collected from four plots across three different depths (0-5 cm, 35- 40 cm, and 95-100 cm) using a Russian auger along a transect each from burnt (3 years post-fire) and intact areas of a tropical peat swamp forest in Brunei. Plots were established in June 2017 as part of a previous study which investigated the effects of fire on carbon dynamics. Detailed descriptions about the site, transect setup, and long-term data on peat physicochemical conditions and carbon emissions have been reported in Lupascu et al (2020). Samples were stored and transported to the laboratory on ice, and were transferred to deep freezers until DNA extraction. Peat porewater physicochemical properties such as water temperature, Electrical Conductivity (EC), Total Dissolved Solids (TDS), salinity, and Dissolved Oxygen (DO) were determined prior to sample collection using previously described methods(7).

### DNA extraction and 16S rRNA marker gene sequencing

Genomic DNA was extracted from all peat samples using the ZymoBIOMICS DNA miniprep kit (Zymo Research, Irvine, CA, USA). Sequence libraries were prepared by amplifying the V4-V5 regions of the 16S rRNA gene from the extracted DNA using the 515F (5’-GTGYCAGCMGCCGCGGTAA-3’) and 926R (5’-CCGYCAATTYMTTTRAGTTT-3’) primers tagged with Illumina overhang adapters. Libraries were sequenced on the Illumina MiSeq (Illumina, San Diego, CA, USA) with 2x 300 bp chemistry at SCELSE (https://www.scelse.sg), Nanyang Technological University, Singapore. We generated a total of 8.7M paired-end reads, with each sample containing on an average 80,105 reads.

### Sequence analyses

Raw sequence data was processed using Cutadapt v3.4 (33) to remove Illumina adapters and phiX sequences. Adapter-free reads were trimmed (forward: 230 bp, reverse: 180 bp) and filtered to retain only those with a maximum error rate (maxEE) ≤3. Overall, 6.1M reads (approx. 70%) were retained post-filtering. Amplicon sequence variants (ASVs) were identified from filtered reads using the DADA2 (34) pipeline. Chimera-free ASVs were filtered to retain only those with length 364 ±10 bp. ASV counts were estimated by mapping filtered reads to length-filtered chimera-free ASVs. Taxonomic labels were inferred using the SILVA database v138.1 (35). The R script used to run this analysis is available on figshare as “btp_16SrRNA_gene_sequence_analysis.R”.

### Microbiome diversity and compositional analysis

Compositional differences between fire-affected and intact peat samples, and across depth were analysed using R v4.2.2 (36) and PRIMER-E v7 (https://www.primer-e.com/). Bray-Curtis dissimilarity between samples were computed using Hellinger-transformed ASV counts. Between-sample variability due to fire and depth were quantified using permutational analysis of variance (PERMANOVA) implemented in PRIMER-E v7 and visualised using non-metric multidimensional scaling (NMDS) routine available in the phyloseq v1.42.0 (37) and vegan v2.6.4 (38) R packages.

Datasets for all analysis were cleaned and wrangled using the tidyverse v1.3.2 (39) R package. Archaea:Bacteria ratio was computed using total sum scaled (TSS) transformed ASV counts. Within-sample diversity was estimated using the Shannon diversity metric with raw counts. The effects of fire on Archaea:Bacteria ratio and diversity was tested using ANOVA. Post-hoc tests were conducted using the emmeans R package; p-values were adjusted using the mvt method. The harmonic mean of the global relative abundance and global prevalence of each ASV was used to identify taxa that are consistently dominant in burnt and intact communities. The magnitude of change in an ASVs relative abundance with fire and depth was quantified using differential abundance analysis implemented in the DESeq2 v1.38.3 (40) R package. Log_2_ fold changes were shrunk using the apeglm v1.20.0 (41) R package for ASVs with either low information across samples or high dispersion; p-values were adjusted for multiple testing. Significantly different taxa were identified as those with an adjusted p-value <0.05.

### Identification of putative methanogens

Putative methanogenic archaea were identified as those archaeal ASVs with a predicted methanogenesis KEGG (42) module(s) (module completeness ≥75%). KEGG modules for each ASV were predicted using the PICRUST2 (43) pipeline. Functional predictions were only retained for those ASVs with a Nearest Sequenced Taxon Index (NSTI) value ≤2. Module completeness was computed using enrichM v0.6.5 (https://github.com/geronimp/enrichM). Putative methanogens identified were verified by cross-referencing with known named groups (e.g. Methanobacteriota) when available.

### Estimation of clade-and community-level assembly processes

Clade-and community-level assembly processes were estimated using a bin-based null modelling approach implemented in the iCAMP v1.5.12 (32) R package. Significant phylogenetic signal is required for estimating assembly processes. A phylogenetic tree was built using FastTree2 (44) with ASV sequences aligned using the DECIPHER (45) R package. Phylogenetic signal was estimated by regressing between-ASV phylogenetic distances against between-ASV niche differences in a pairwise manner and tested for statistical significance using a Mantel test (Mantel R ≥0.1, P<0.05). The distance within which phylogenetic signal was consistently observed was used as the phylogenetic distance threshold for demarcating phylogenetic bins.

The beta net relatedness index (bNRI) based on the beta mean pairwise distance (bMPD) was computed within each bin and then compared against those computed under null expectations to estimate the relative contribution of homogeneous (bNRI >- 1.96) and heterogenous selection (bNRI <+1.96) to microbiome assembly. Relative contributions of stochastic processes to microbiome assembly were estimated when |bNRI| was ≤1.96, by computing and thresholding the modified Raup-Crick (RC) metric (Homogenizing dispersal RC <-0.95, dispersal limitation RC >+0.95, drift otherwise). Finally, relative contributions of different selective and stochastic processes at the community-level were estimated using an abundance-weighted approach that takes into consideration the relative abundance of each ASV within the community.

The effect of fire on the assembly of entire communities and individual clades were tested by computing standardised effect sizes (Cohen’s d) and tested using a one-sided bootstrap test (1000 times). Cohen’s d was computed as the difference in means between the fire and intact samples divided by the combined standard deviation. Computed effect sizes were classified as large (|d| >0.8), medium (0.5 < |d| ≤ 0.8), small (0.2 < |d| ≤ 0.5), or negligible (|d| ≤ 0.2).

### Relationships between assembly processes and physicochemical conditions

Correlations between the relative importance of assembly processes and environmental variables were tested using Mantel tests implemented in the iCAMP (32) R package. We computed associations between pairwise differences in the relative importance of each assembly process with either the variation (absolute difference) or mean of every environmental variable. Mantel tests with different types of data-transformations (untransformed or log-transformed) were tested in order to identify the best model. All environmental variables were log-transformed when required except pH.

## Results

### Peat physicochemical conditions remain strongly altered four years post-fire

Peat samples were collected from four points across three different depths (0-5 cm, 35-40 cm, and 95-100 cm) along a transect each in the fire-affected and intact areas of tropical peat dome in Brunei. We analysed peat porewater from these three depths to examine if physicochemical conditions remained altered four years post-fire. Water temperature (+0.4°C, ANOVA, *p*=0.006; Supplementary Figure 1) and pH (+0.56, ANOVA, *p*=0.007; Supplementary Figure 1) remained significantly elevated in burnt peat compared to intact peat. There were no significant differences in pH across depths (ANOVA, *p*=0.16) but water temperature was significantly higher (Pairwise t-test, *p*<0.001 for all comparisons) at the surface. On the other hand, although EC (ANOVA, *p*=0.07), TDS (ANOVA, *p*=0.07), and salinity (ANOVA, *p*=0.07) were lower in burnt peat (Supplementary Figure 1), they did not meet our thresholds for statistical significance presumably due to the lower number of samples collected in this study. Porewater oxygen concentrations did not significantly differ between burnt and intact peat (ANOVA, *p*=0.60) but showed a decreasing trend with depth (Supplementary Figure 1) that approached statistical significance (ANOVA, *p*=0.08). Overall, except for dissolved oxygen concentrations, these results are consistent with those reported earlier (7) from the same site and show that fire-induced changes in peat physicochemical conditions continue to persist four years post-fire.

### Fire increases relative abundance of Archaea in deep peat and shifts community composition

To study the effects of fire on tropical peat microbiomes, we analysed bacterial and archaeal communities using 16S rRNA marker gene sequencing. Overall, we detected 3,686 bacterial and 242 archaeal ASVs. Fire significantly reduced bacterial dominance in deep peat (Pairwise t-test, *p*<0.001) with archaea becoming increasingly abundant in the community (Figure 1a). However, archaeal diversity did not differ significantly with either fire (ANOVA, *p*>0.99) or depth (ANOVA, *p*=0.42) (Figure 1b). In contrast, fire significantly decreased bacterial diversity only in deep peat (Pairwise t-test, *p*=0.006) (Figure 1c).

**Figure 1.**
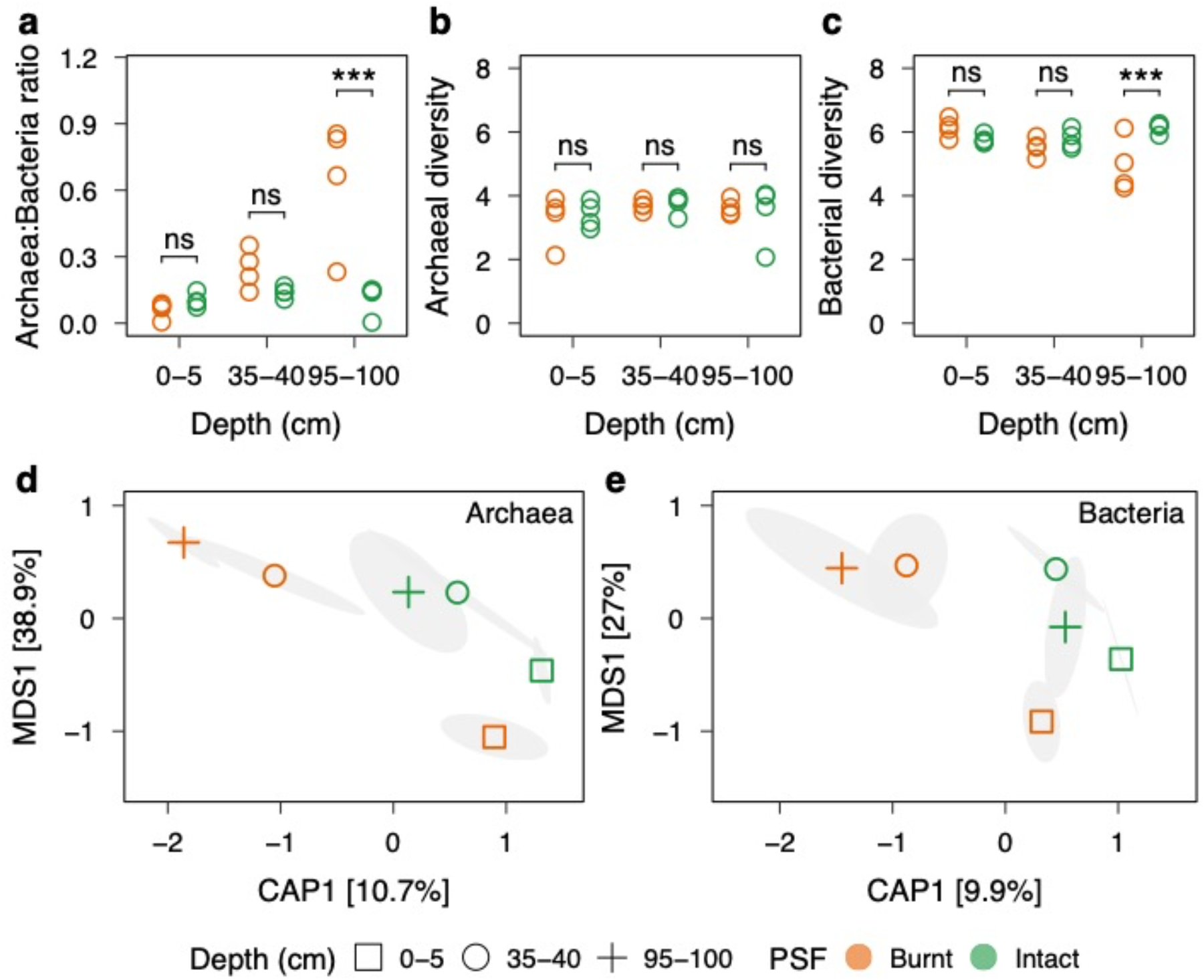
Fire strongly alters microbiome diversity and composition. (a) Fire significantly increases Archaea:Bacteria ratio in deep peat (Pairwise t-test, *p*<0.001), (b) Archaeal diversity does not significantly differ with fire (ANOVA, *p*>0.99) or depth (ANOVA, *p*=0.42), (c) Fire reduces bacterial diversity in deep peat (Pairwise t-test, *p*=0.006) (d) and (e) Fire significantly shifts both archaeal and bacterial communities across all depths. Symbols represent centroids with 95% confidence ellipses. Figure 2. Shifts in archaeal and bacterial communities at the class level. Top five classes were identified based on the harmonic mean of relative abundance and prevalence. Together, these classes account for approximately 68.8-81.64% of bacterial and 98.28-99.53% of archaeal communities respectively.

As opposed to the limited changes in diversity, composition of both archaeal and bacterial communities were significantly different between burnt and intact peat (Figure 2d) (archaea PERMANOVA, *p*=0.003; bacteria PERMANOVA, *p*=0.004; Supplementary Table 1). Although, shifts in community composition were observed across all three depths (archaea PERMANOVA, *p*=0.001; bacteria PERMANOVA, *p*=0.006; Supplementary Table 1), the water-saturated peat layers (35-40 cm and 95- 100 cm) were impacted to a greater extent (Figure 1d and 1e). Depth-stratification was, however, similar across both burnt and intact peat; communities from the surface were significantly different to those from the middle (Pairwise t-test, Archaea *p*=0.001, Bacteria *p*=0.005) and deeper (Pairwise t-test, Archaea *p*=0.001, Bacteria *p*=0.03) layers. Together, these results show that fires shift both archaeal and bacterial composition across depth and is linked to increased susceptibility of bacteria in deep peat resulting in less diverse bacterial communities. As opposed to bacteria, archaea appear to be more resistant to changes post-fire.

**Figure 2.**
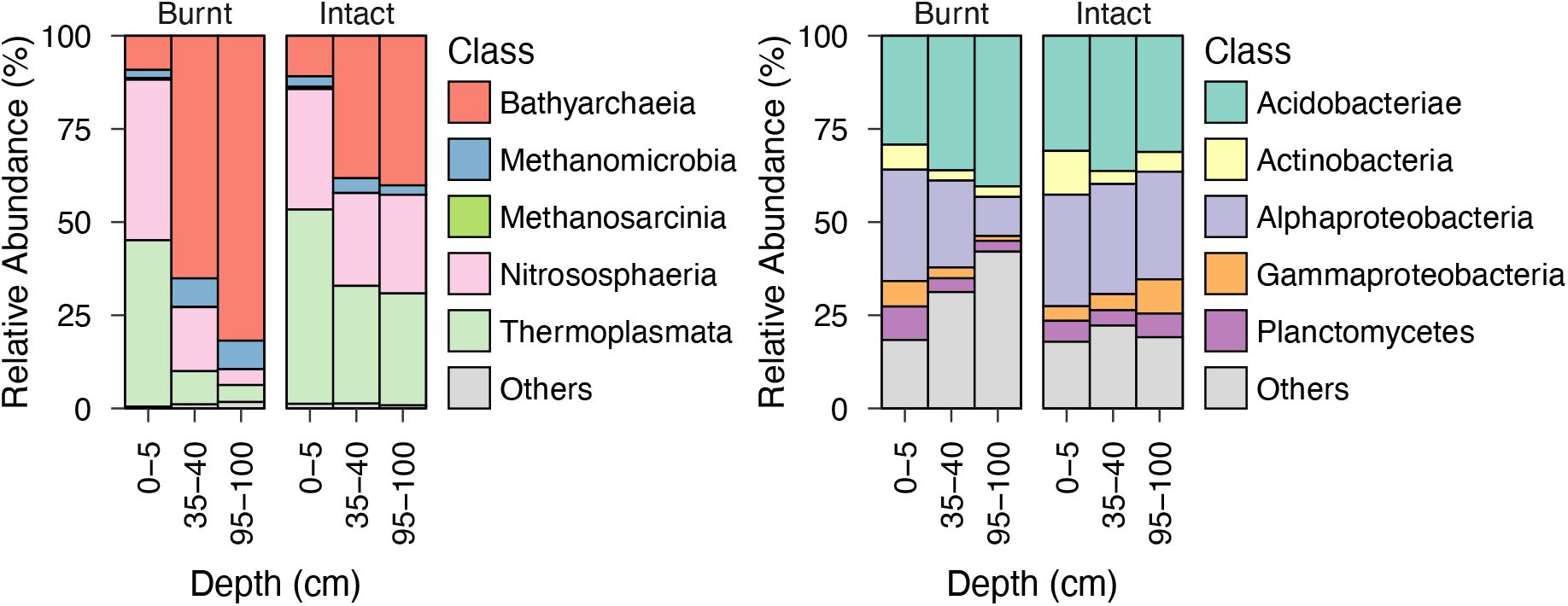
Shifts in archaeal and bacterial communities at the class level. Top five classes were identified based on the harmonic mean of relative abundance and prevalence. Together, these classes account for approximately 68.8-81.64% of bacterial and 98.28-99.53% of archaeal communities respectively.

### Bathyarchaeia respond strongly to fires

Based on the prevalence and abundance of bacterial and archaeal taxa across all samples, we identified those that dominate communities and respond significantly to fires. Fire-induced compositional shifts were apparent even at broad taxonomic levels examined here, particularly in below-surface peat (35-40 cm and 95-100 cm). At depths ≥30 cm, Acidobacteriae were dominant in both burnt (approx. 35.26%) and intact peat (approx. 32.80%) (Figure 2a). Altered peat conditions at depths ≥95 cm significantly enriched for several bacterial classes such as Dehalococcoidia (Wald’s test, *p*<0.001), Methylomirabilia (Wald’s test, *p*=0.01), and Spirochaetia (Wald’s test, *p*=0.002), whereas it significantly decreased the abundance of classes such as the Gammaproteobacteria (Wald’s test, *p*=0.046) (Supplementary Table 2). Archaeal communities at depths ≥30 cm was dominated by the Bathyarchaeia, which nearly doubled in abundance in burnt peat compared to intact peat (Wald’s test, *p*=0.03) (Figure 2b). Fire also significantly increased the abundance of archaeal classes such as Methanomicrobia (Wald’s test, *p*=0.01) and Methanomethylicia (Wald’s test, *p*=0.01) at depths ≥95 cm (Supplementary Table 2). In contrast to communities at depths ≥30 cm, profiles of abundant taxa in surficial peat were qualitatively similar between burnt and intact peat. Bacterial communities in surface peat were dominated by Acidobacteriae and Alphaproteobacteria, whereas archaeal communities were dominated by Thermoplasmata and Nitrososphaeria.

### Fires increase the relative abundance of diverse methanogenic archaea

Fire significantly increases methane emissions (7), production of which is constrained to a group of strictly anaerobic methanogenic archaea. To determine the impact of fire on methanogens, we identified putative taxa capable of generating methane, methanogenesis pathways, and estimated their relative abundance in burnt and intact peat. We identified 37 archaeal taxa from seven different classes capable of generating methane through hydrogenotrophic, acetoclastic, and methylotrophic pathways (Supplementary Table 3). Fire significantly increased the abundance of methanogens only at peat depths ≥95 cm (ANOVA, *p*<0.001; Supplementary Figure 2). This was driven to a large extent by increases in the relative abundance of methanogens from the classes Bathyarchaeia (approx. 100x) and Methanomicrobia (approx. 2x) respectively compared to intact peat. Pathway predictions showed that while both classes were capable of generating methane through acetoclastic methanogenesis, they differ in their capacity to generate methane from carbon-dioxide (Methanomicrobia) and C1 compounds (Bathyarchaeia) respectively. Although, fire significantly increased the abundance of methanogenic archaea at depths ≥95 cm, it did not significantly impact their diversity (ANOVA, *p*=0.08). Together, these results show that diverse methanogenic archaea can persist or increase in abundance in altered peat conditions post-fire at depths ≥95 cm. These conditions may further favour the growth of methylotrophic methanogens such as those from the class Bathyarchaeia.

### Homogeneous selection shapes archaeal communities post-fire

We next investigated if archaeal communities and clades containing putative methanogens shift in a predictable manner post-fire by determining the relative importance of different deterministic and stochastic assembly processes at the community-and clade-level. Deep peat (depth ≥95 cm) archaeal communities which showed the strongest response post-fire were binned into six clades with significant phylogenetic signal (Supplementary Table 4). Fires imposed strong, consistent selective pressures resulting in highly homogenised archaeal communities (Cohen’s d: 4.5, *p*<0.001; Supplementary Table 5) (Figure 3). This result was consistent with analysis of multivariate dispersions which showed that communities in burnt peat were significantly more similar to each other compared to those in intact peat (PERMDISP, *p*=0.03). On the other hand, fire significantly decreased the relative importance of ecological drift (1.3-12% turnovers; Cohen’s d: −2.6, *p*=0.02; Supplementary Table 5) compared to intact peat (10-82% turnovers).

**Figure 3.**
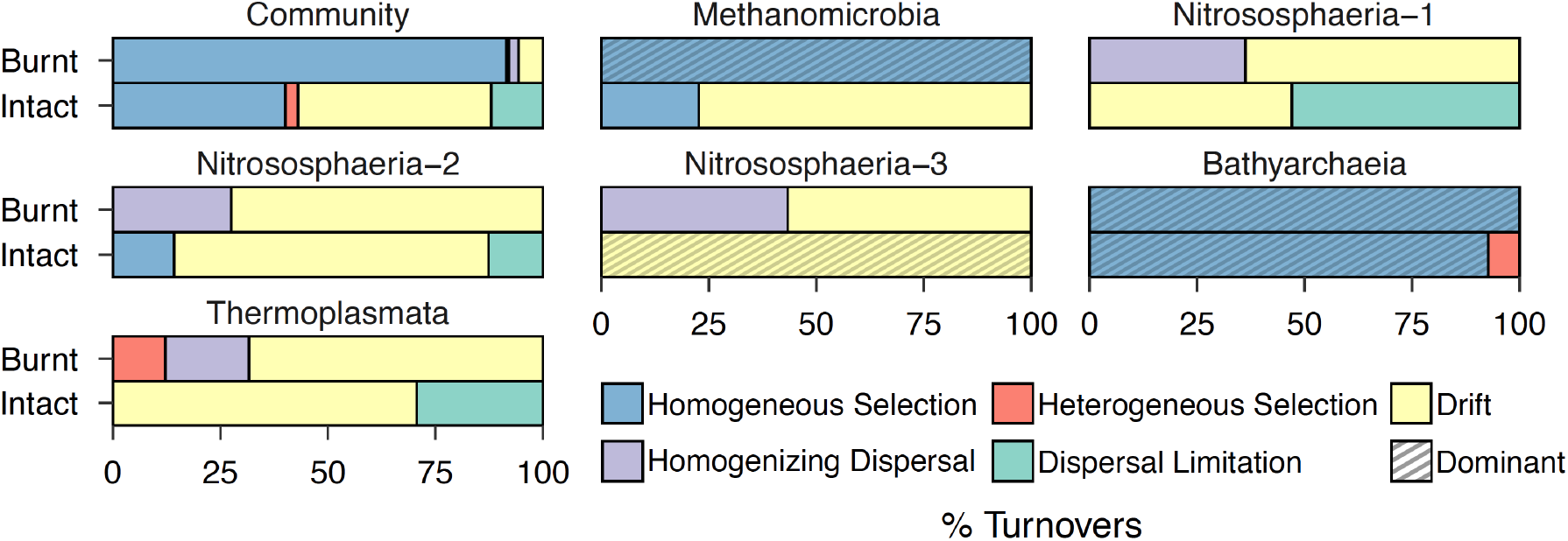
Homogeneous selection assemble archaeal communities post-fire. Fire shifts relative importance of assembly processes both at the community and clade-level. Dominant process acting on each clade is indicated with dashed lines.

Bathyarchaeia (83.9%) and Methanomicrobia (7.5%), which include abundant methanogens, contributed most to the increase in homogenous selection post-fire (Figure 3; Supplementary Table 6). Fire significantly increased the relative importance of homogeneous selection in assembling Methanomicrobia (One-sided bootstrap test, *p*<0.001; Supplementary Table 7), however, it did not alter the relative importance of processes shaping the Bathyarchaeia, which were predominantly assembled by homogeneous selection in both burnt and intact peat. In contrast to the Bathyarchaeia and Methanomicrobia, clades consisting of the Nitrososphaeria and Thermoplasmata were assembled to a large extent by stochastic processes in both burnt and intact peat. Collectively, these results show that fires select for phylogenetically clustered communities driven largely by the classes Bathyarchaeia and Methanomicrobia.

### Homogeneous selection and drift shift with porewater DO concentrations

Finally, we determined the influence of porewater physicochemical conditions on the relative importance of different assembly processes using Mantel tests. We focussed only on homogeneous selection (91.5% in burnt and 40.1% in intact peat) and drift (5.7% in burnt and 44.9% in intact peat) as they accounted for a major proportion of turnovers in both burnt and intact peat. Mantel tests showed that homogenous selection increased significantly with decreasing concentrations of porewater dissolved oxygen (mean DO, Pearson’s R: −0.8, *p*=0.01; Figure 4) and EC (mean EC, Pearson’s R: −0.6, *p*=0.02; Figure 4). The strong negative correlation between homogenous selection and DO remained significant when controlling for other factors. In contrast, ecological drift increased significantly with increasing concentrations of porewater DO (mean DO, Pearson’s R: 0.6, *p*=0.01; Figure 4) and this relationship was preserved when other factors were controlled for. These results show that porewater dissolved oxygen concentration may play an important role in determining the relative importance of homogeneous selection and drift.

**Figure 4.**
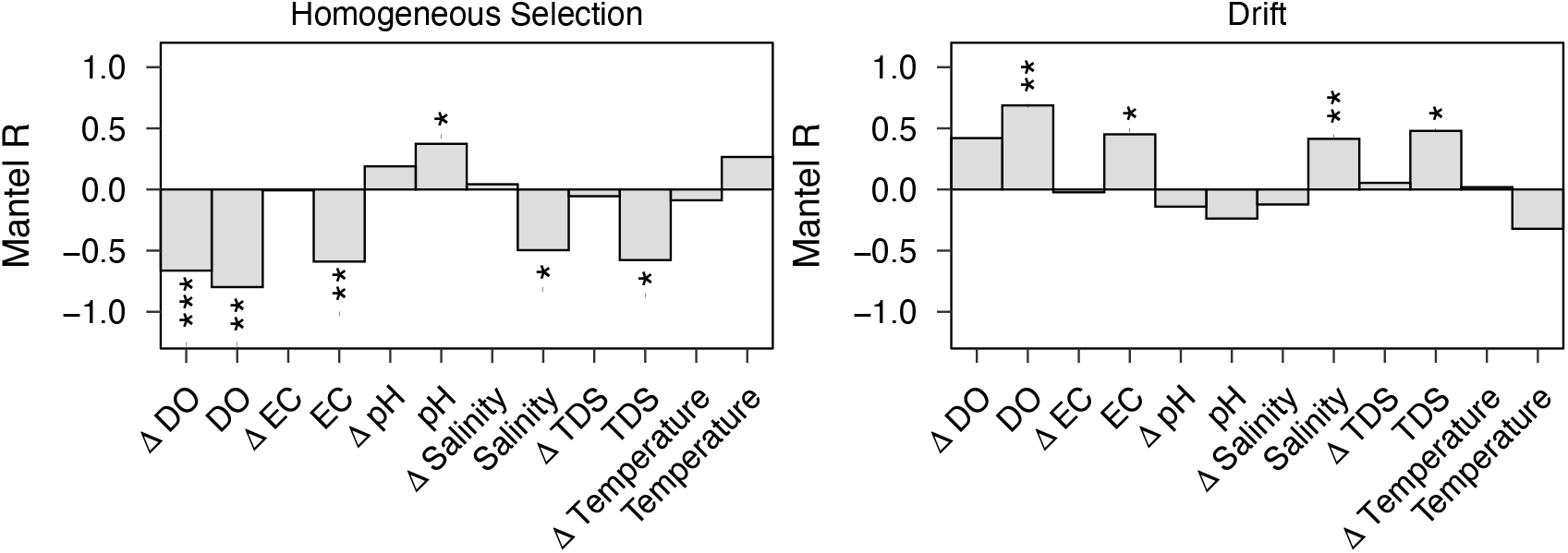
Homogeneous selection and drift shift with porewater DO concentrations. Associations between homogeneous selection, drift, and porewater physicochemical conditions based on Mantel test. Significance is expressed as *** *p*<0.01; ** *p*<0.05; * *p*<0.1

## Discussion

Fires have long-lasting impacts on the carbon dynamics of tropical peatlands, yet we lack an understanding of its impacts on peat microbiome composition and functioning. Here, we not only show for the first time that fire alters diversity and composition of tropical peat microbiomes, but also its predictability based on environmental conditions. We also found that specific methanogens respond positively to fires in a predictable manner thereby reconciling altered physicochemical conditions with elevated methane emissions post-fire observed earlier. Overall, this study highlights the complex interplay between fire, environmental factors, and microbiomes in shaping the functioning of burnt tropical peatlands. Furthermore, our results provide important insights into microbiome assembly that can guide efforts to engineer the peat microbiome to restore ecosystem services post-fire.

Fire had strong but varying effects on the relative abundance and diversity of bacteria and archaea. Our study confirms earlier reports that fires reduce bacterial abundance and diversity. However, we observed these effects at depths >1m below the peat surface in contrast to earlier studies which report such strong changes mainly at the soil/peat surface (20, 22). Therefore, we conclude that fires can have long-lasting impacts on bacterial diversity especially in deeper peat layers. Our earlier study showed little to no differences in peat characteristics such as bulk density, C/N ratio and peat age at these depths (7) which suggests negligible direct effects of burning on bacterial diversity and abundance, although loss due to an increase in temperature immediately post-fire cannot be ruled out. Conversely, significantly lower EC and DO which indicate limited availability of energetically-favourable Terminal Electron Acceptors (TEA) likely explain the reduction in bacterial diversity and abundance to a greater extent, similar to low-energy deeper layers in highly-stratified environments (for example, lake sediments) (46, 47). However, higher pH indicates the presence of nutrients albeit in a reduced state due to lower DO, which in turn was attributed to prolonged periods of flooding (7) and a lack of root oxygenation at these depths associated with a switch from deep-rooted trees to shallow-rooted graminoids (48, 49). This strongly suggests an overarching role of aboveground vegetation on the belowground redox landscape and consequently community turnover. Several studies indeed have shown that secondary plant succession post-fire can strongly influence belowground microbiome diversity and abundance (27, 50, 51). On the other hand, our findings on archaeal diversity agrees with an increasing number of studies which either show no difference or recovery to pre-fire levels 1-2 years post-fire (52, 53). An increase in their relative abundance at these depths is consistent with their capacity to persist and grow in low-energy environments (46, 47) such as those encountered post-fire described above. Although we observed significant changes in relative abundance, we recognise that this may not necessarily reflect a change in absolute abundance. Future studies must therefore determine changes in absolute abundance as they can provide more detailed insights into ecological strategies (for example, increased growth versus reduced competition for resources) post-fire.

Our results indicate that the effects of fire on community composition are more widespread across different depths compared to its impact on diversity, an effect which has been reported in previous work. Changes in community structure mirror significant differences in physiochemical conditions which continue to persist four years post fire. In particular, changes in pH and temperature post-fire can extensively restructure microbiomes, effects of which are well established across numerous ecosystems (54, 55, 56). Differences in community structure across depth in both burnt and intact peat reflect changes in water table levels and consequently oxygen availability, an effect that has been extensively documented in peatland ecosystems (57, 58). Additionally, frequent flooding post-fire resulting from peat subsidence may further accentuate depth-specific differences in community structure. Furthermore, the functional consequences of fires disrupting the depth-stratification in community structure merits further investigation as important vertically-structured source-sink relationships (58) (for example, methane production and consumption) can be significantly altered post-disturbance (59).

The response of specific taxa to fire at depths ≥1m further support our assertion that factors such as the limited availability of preferred terminal electron acceptors (for example, oxygen) driven by changes in aboveground vegetation play a greater role in determining community composition at these depths. For example, Bathyarchaeia (60) and Dehalococcoidia (61) which thrive in anoxic low-energy environments increased in relative abundance whereas Gammaproteobacteria which are generally copiotrophic in nature (62) decreased in relative abundance. In particular, the remarkable increase in the relative abundance of Bathyarchaeia post-fire is consistent with previous reports (52) which show that they can respire refractory carbon and are particularly dominant in anoxic niches with fluctuating redox regimes (60). It is important to note that our conclusion remains unchanged even though it is possible that the observed increase in relative abundances may be due to a decrease in the absolute abundances of other taxa. Our previous study revealed the existence of millennia-old carbon at these depths (7) and hence a shift in metabolic strategies from labile to refractory carbon through mechanisms such as priming likely has important implications for the turnover of old carbon.

A significant increase in the abundance of methanogenic archaea at depths ≥1m in burnt peat corroborates earlier work at our site that found elevated methane flux 2-3 years post-fire. Methane flux in burnt peat was particularly high during prolonged periods of flooding but remained comparable to those from intact peat otherwise (7). This shows that the redox regime and consequently the availability of different TEAs fluctuates at these depths, depletion of which precedes methanogenesis. Putative methanogens from the classes Methanomicrobia and Bathyarchaeia contributed most to this increase in burnt peat, an observation consistent with previous reports which show that Methanomicrobia increase in relative abundance in high methane-flux habitats (63). On the other hand, a rather unique finding from our study is the remarkable increase in the relative abundance of putative methanogens from the class Bathyarchaeia, which suggests that they likely play a crucial role in methane emissions from fire-affected tropical peatlands. Further, our findings are consistent with studies which show a positive co-abundance relationship between Methanomicrobia and Bathyarchaeia (60, 64) which either suggests metabolic interconnectedness or that they occupy a similar niche. Functional predictions further showed that putative methanogens from the class Bathyarchaeia are capable of both acetoclastic and methylotrophic methanogenesis, which likely offers them a competitive advantage in burnt peat where there is increased availability of plant-derived labile carbon especially during the wet season. However, we acknowledge that these observations are based on predicted functions and therefore need additional validation with metagenomic and metatranscriptomic studies.

Consistent with our hypothesis, the relative importance of homogeneous selection strongly increased post-fire at depths ≥1m which shows that changes in archaeal community composition can be predicted based on environmental conditions and functional traits. Our findings agree with previous studies which have shown that the relative importance of homogeneous selection increases with time since fire and can account for a majority of community turnovers 3-4 years post-fire (27, 28, 65). Consistent selective pressures on local communities can arise from either increased environmental homogeneity or stressful environmental conditions. In contrast to previous studies (27), we did not observe decreased variability in environmental conditions post-fire, despite observing greater homogeneity in aboveground plant communities and microtopography (7, 48). Instead, our finding that homogenous selection is inversely correlated with porewater DO concentrations and salinity suggests that the limited availability or absence of energetically favourable TEAs impose strong selective pressures which homogenize the community. Indeed, it has been shown that low-energy environments exert strong selective pressures either by favouring taxa that can tolerate stressful conditions or by increased competition for scarce resources (46, 66). Our finding that stochasticity plays a crucial role in shaping archaeal communities in intact peat is consistent with prior work that has demonstrated its significance in productive environments (67). Overall, our findings indicate that archaeal communities have not fully recovered to pre-fire states due to altered physicochemical conditions which continue to persist four years post-fire.

Clade-specific differences in assembly processes post-fire is a rather unique finding as most studies only estimate such processes at the community level. The contrasting responses of Methanomicrobia and Bathyarchaeia post-fire likely reflect differences in both physiology and members comprising the metacommunity. The strong selection of Bathyarchaeial lineages in intact peat indicates the presence of highly-adapted taxa in the seed pool, whose assembly remains largely unperturbed post-fire likely due to their metabolic versatility. Indeed, previous studies have demonstrated that metabolic generalists are more resilient to environmental perturbations (68). In contrast, the primary mechanism shaping the Methanomicrobia shifted from ecological drift to homogeneous selection post-fire likely driven by competition for substrates such as acetate and hydrogen in a low-energy environment. For instance, prior work has shown that in energy-limited sediments, sulfate reducing bacteria routinely outcompete methanogens for these so-called competitive substrates (69). Thus, our results suggest that the composition of methanogenic clades can be predicted based on environmental conditions four-years post-fire, mechanisms driving which can be elucidated in the future using substrate-addition experiments.

An important limitation is our findings represent a snapshot in time, despite being informed by long-term monitoring of specific ecosystem functions (7). Microbiome functioning likely changes with seasonal changes in precipitation as evinced by temporal variability in carbon dioxide and methane emissions. This underscores the need for long-term studies which capture successional changes in microbiome composition and functioning post-fire. Although we cross-referenced functional predictions with existing literature when available, our interpretations about functional potential should be treated with caution and validated using approaches which measure activity, due to known limitations with predictive approaches. Finally, future studies must consider bacterial and fungal contributions to changes in ecosystem functioning post-fire and their influence on archaeal communities. Despite these limitations, our findings highlight important ecological principles that could be used to disrupt functioning of non-beneficial clades and engineer microbiomes that can assist ecosystem recovery post-fire. For instance, efforts to disrupt the activity of methanogens should focus on re-establishing the native plant community. On the other hand, the negligible role of dispersal limitation together with the overarching role of homogeneous selection post-fire suggests that efforts to assist plant establishment should focus on rhizosphere inoculation strategies with locally-sourced beneficial microorganisms which can withstand the harsh post-fire environment.

We show that environmental conditions and peat microbiomes remain deeply affected four-years post-fire. Our results indicate that the harsh post-fire environment characterised by fluctuating redox conditions and limited availability of energetically favourable TEAs especially at depths ≥1m drive reductions in bacterial diversity and changes in microbiome composition. Such a low-energy environment in turn enriches for diverse methanogenic archaea, a finding which corroborates earlier work which showed elevate methane flux 2-3 years post-fire. Finally, our results shows that changes in archaeal composition and clades containing abundant methanogens can be predicted based on environmental conditions and functional traits. Collectively, our results demonstrate that peat microbiomes follow predictable trajectories post-fire based on macroecological principles with key management implications for post-fire ecosystem recovery.

## Author contributions

HA is a biogeochemist while AB is a microbial ecologist; both contributed equally to this manuscript. HA and ML designed the study and collected samples. HA processed the samples. AB analysed the data, generated the figures and tables. AB and HA wrote the manuscript with critical inputs from ML and SS.

## Acknowledgements

We would like to thank the Ministry of Education, Singapore and the Integrated Tropical Peatlands Research Programme (INTPREP) hosted at the NUS Environmental Research Institute (NERI) for funding this research; the Brunei Forestry Department for entry and sampling permits (Ref:[209]/JPH/UND/17 PT.1); the Biodiversity Research and Innovation Centre for export permits; the Biosecurity

Division for phytosanitary certificates; Universiti Brunei Darussalam for permission to conduct research; Wardah Haji Tuah, Hazimah Haji Mohammad Din and Salwana Md Jaafar for logistics and field support; Mohammad Azizi bin Mohammad Firdaus Chee, Dk Noor Ummiatul Aiqah binti Pg Zainalabidin, Siti Nur Aqilah Afizah binti Mohd. Alias, Rachel Wee, Vidya Veloo for their help in the field.

## Data availability

Raw sequence data is available on the NCBI Sequence Read Archive (SRA). Datasets and data products generated from the raw data are available on figshare.

## Code availability

R codes used to analyse the data are available on GitHub.

