## Supplementary Information for "High methane flux in a tropical peatland post-fire is linked to homogenous selection of diverse methanogenic archaea"

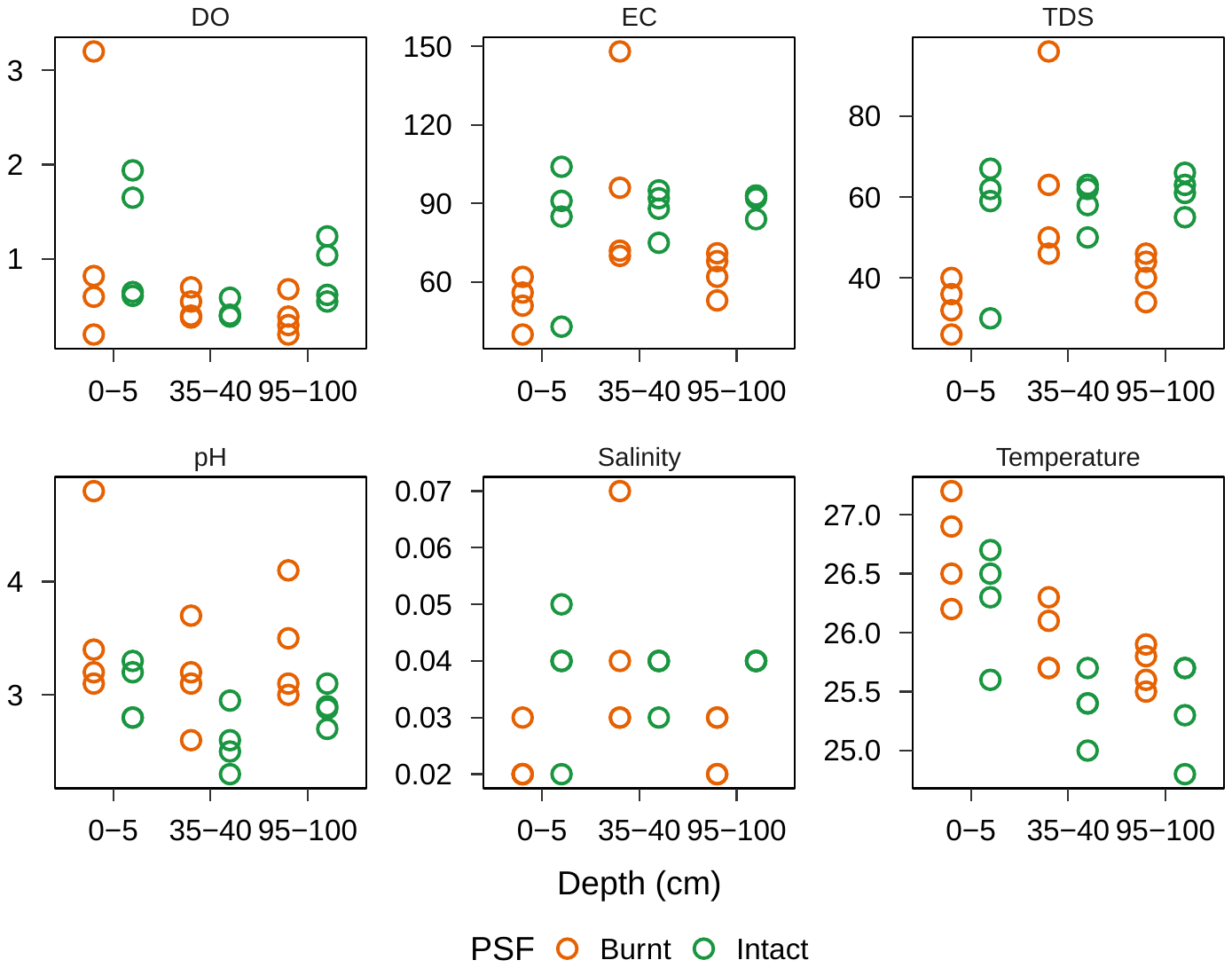

**Supplementary Figure 1:** Peat porewater physicochemical conditions remain strongly altered four-years post-fire.

|  | Factor | F-value | P-value | R^2^ |
| --- | --- | --- | --- | --- |
| Archaea | Fire | 3.79 | 0.003 | 18.43 |
|  | Depth | 5.34 | 0.001 | 28.15 |
|  | Fire:Depth | 1.27 | 0.247 | 9.98 |
| Bacteria | Fire | 2.88 | 0.004 | 16.38 |
|  | Depth | 2.85 | 0.006 | 19.92 |
|  | Fire:Depth | 1.31 | 0.18 | 11.62 |

**Supplementary Table 1:** PERMANOVA summary statistics using Type III Sums of Squares. Fire and depth were considered as fixed factors. P-values were estimated based on 999 permutations.

**Supplementary Table 2:** Significantly enriched or depleted bacterial and archaeal classes in burnt peat at depth ≥95 cm. Table is provided in a separate spreadsheet.

**Supplementary Table 3:** List of putative methanogens. Table is provided in a separate spreadsheet.

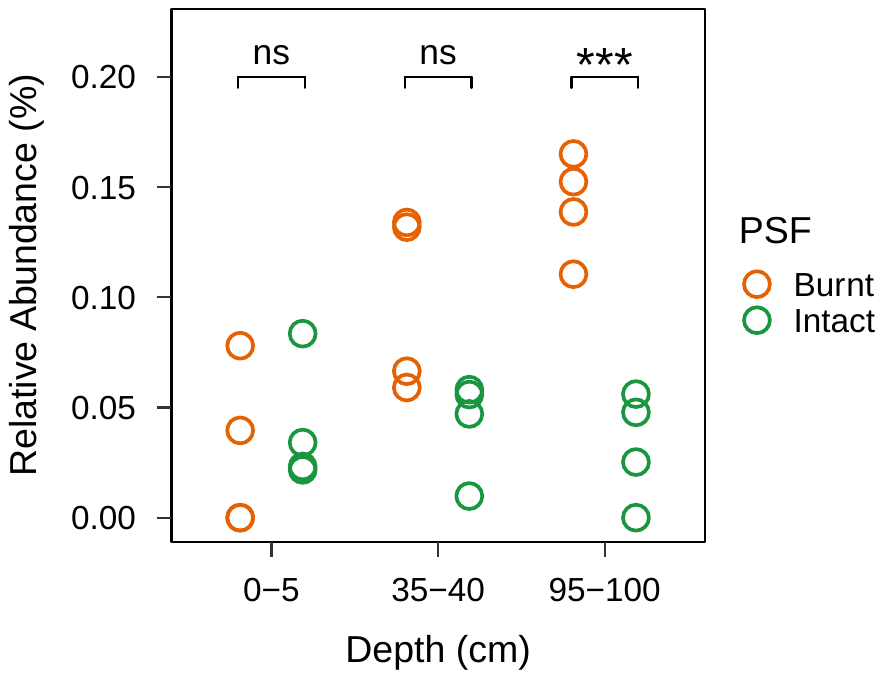

**Supplementary Figure 2:** Methanogens are significantly more abundant at 1m depth post-fire.

**Supplementary Table 4:** ASVs comprising each phylogenetic bin. Table is provided in a separate spreadsheet.

**Supplementary Table 5:** Relative importance of different ecological processes structuring Archaeal communities in burnt and intact peat. Values for burnt and intact peat show observed values (with standard deviations). Statistical significance was estimated using a one-sided bootstrap test.

| Process | Burnt | Intact | d | Effect size | P |
| --- | --- | --- | --- | --- | --- |
| Homogeneous selection | 91.47 (0.02) | 40.1 (16.26) | +4.45 |  | <0.001 |
| Heterogeneous selection | 0.54 (0.83) | 2.94 (4.54) | -0.76 | Large | 0.56 |
| Ecological drift | 5.66 (3.9) | 44.93 (21.35) | -2.56 | Large | 0.02 |
| Homogenizing Dispersal | 2.32 (1.76) | 0 | +1.86 |  | 0.11 |
| Dispersal limitation | 0 | 12.04 (11.67) | -1.48 |  | 0.35 |

**Supplementary Table 6:** Contribution of processes shaping each clade to ecological processes at the community level.

|  |  | Phylogenetic Bins | | | | | |
| --- | --- | --- | --- | --- | --- | --- | --- |
| Fire | Process | 1 | 2 | 3 | 4 | 5 | 6 |
| Burnt | Heterogeneous Selection | 0% | 0% | 0% | 0% | 0% | 0.54% |
|  | Homogeneous Selection | 7.6% | 0% | 0% | 0% | 83.9% | 0% |
|  | Dispersal Limitation | 0% | 0% | 0% | 0% | 0% | 0% |
|  | Homogenizing Dispersal | 0% | 0.6% | 0.3% | 0.5% | 0% | 0.9% |
|  | Drift | 0% | 1% | 0.9% | 0.7% | 0% | 3% |
| Intact | Heterogeneous Selection | 0% | 0% | 0% | 0% | 2.9% | 0% |
|  | Homogeneous Selection | 0.7% | 0% | 1.7% | 0% | 37.7% | 0% |
|  | Dispersal Limitation | 0% | 2.5% | 1.5% | 0% | 0% | 8% |
|  | Homogenizing Dispersal | 0% | 0% | 0% | 0% | 0% | 0% |
|  | Drift | 2.4% | 2.2% | 8.9% | 12.1% | 0% | 19.4% |

**Supplementary Table 7:** Fire-induced changes in the relative importance of different ecological processes shaping each clade.

|  |  | Phylogenetic Bins | | | | | |
| --- | --- | --- | --- | --- | --- | --- | --- |
| Fire | Process | 1 | 2 | 3 | 4 | 5 | 6 |
| Burnt | Heterogeneous Selection | 0% | 0% | 0% | 0% | 0% | 12% |
|  | Homogeneous Selection | **100%** | 0% | 0% | 0% | **100%** | 0% |
|  | Dispersal Limitation | 0% | 0% | 0% | 0% | 0% | 0% |
|  | Homogenizing Dispersal | 0% | 36% | 28% | 43% | 0% | 19% |
|  | Drift | 0% | 64% | 72% | 57% | 0% | 68% |
|  | Process Importance | 100% | 64% | 72% | 57% | 100% | 68% |
|  | P-value | **0.00** | 0.30 | 0.30 | 0.33 | **0.00** | 0.30 |
| Intact | Heterogeneous Selection | 0% | 0% | 0% | 0% | 7% | 0% |
|  | Homogeneous Selection | 23% | 0% | 14% | 0% | **93%** | 0% |
|  | Dispersal Limitation | 0% | 53% | 13% | 0% | 0% | 29% |
|  | Homogenizing Dispersal | 0% | 0% | 0% | 0% | 0% | 0% |
|  | Drift | 77% | 47% | 73% | **100%** | 0% | 71% |
|  | Process Importance | 77% | 53% | 73% | 100% | 93% | 71% |
|  | P-value | 0.14 | 0.45 | 0.26 | **0.00** | **0.05** | 0.26 |
